# Shifting patterns of seasonal influenza epidemics

**DOI:** 10.1101/268060

**Authors:** Pietro Coletti, Chiara Poletto, Clément Turbelin, Thierry Blanchon, Vittoria Colizza

**Author notes:** Current address Universiteit Hasselt, I-Biostat, 3500 Hasselt, Belgium (Pietro Coletti).

## Abstract

Seasonal waves of influenza display a complex spatiotemporal pattern resulting from the interplay of biological, socio-demographic, and environmental factors. At country level many studies characterized the robust properties of annual epidemics, depicting a typical season. Here we analyzed season-by-season variability, introducing a clustering approach to assess the deviations from typical spreading patterns. The classification is performed on the similarity of temporal configurations of onset and peak times of regional epidemics, based on influenza-like-illness time-series in France from 1984 to 2014. We observed a larger variability in the onset compared to the peak. Two relevant classes of clusters emerge: groups of seasons sharing similar recurrent spreading patterns (clustered seasons) and single seasons displaying unique patterns (monoids). Recurrent patterns exhibit a more pronounced spatial signature than unique patterns. We assessed how seasons shift between these classes from onset to peak depending on epidemiological, environmental, and socio-demographic variables. We found that the spatial dynamics of influenza and its association with commuting, previously observed as a general property of French influenza epidemics, applies only to seasons exhibiting recurrent patterns. The proposed methodology is successful in providing new insights on influenza spread and can be applied to incidence time-series of different countries and different diseases.

## Introduction

Understanding influenza spatial dynamics is highly relevant to improve preparedness and control, as annual influenza epidemics represent a serious burden for public health worldwide [11]. Several empirical and modeling studies have uncovered a set of key features of disease spatial transmission observed across several seasons. Influenza diffuses through complex spatiotemporal patterns that appear to change across scales [62, 61, 32, 15, 31, 17, 21, 23, 39, 65, 49, 22]. A hierarchical spread combining a wave-like behavior on the small scale and long-range seeding events synchronizing distant populations was reported by several studies [62, 61, 17, 65, 50]. This is generally explained by the multiscale nature of mobility patterns of individuals [12], with commuting mainly acting on the short distance whereas air travel flows are responsible for non-local diffusion [12, 61, 17, 22]. The relative importance of wave-like vs. long-range diffusion and of the associated mobility modes however depends on the geographic scale of the country. Long range coupling by air travel was found in previous analyses of epidemiological data in the US [61, 12, 18, 16], though its role is currently debated [22]. A higher synchrony of epidemics is reported in smaller countries, e.g. in Israel [32] and in France [15, 21, 23] where no dominant transportation mode was identified [23]. Spatial patterns of influenza spread also depend on the dominant viral strain circulating [61, 50, 19, 40, 43], with a tendency of B-dominated influenza seasons to be characterized by slower and later epidemics compared to A-dominated epidemics [50, 19].

Seasonality affects influenza spatial dynamics in multiple ways [37]. Day/night cycles are thought to influence the immune system increasing the susceptibility to infection during winter period [26]. Low absolute humidity and temperature may increase virus survival and overall transmissibility [52], facilitate transmission onset [31], and possibly lead to detectable signals in influenza activity [53, 22, 57, 24].

The vast majority of these studies highlight general and robust tendencies of influenza epidemics and the properties of a *typical* season [62, 61, 32, 31, 21, 23, 65, 56, 50]. Evidence of multiple spatial patterns beyond the one associated to a typical seasonal behavior is however available. Marked radial patterns are for example observed in 4 influenza seasons in the US out of the 8 seasons (2002-2010) under study by Charu *et al.* [22], and monotonous spatial patterns (i.e. highly synchronized) are reported in Japan compared to multitonous ones (i.e. multi-seeding followed by radial diffusion) in the period 1992-1999 [49]. While the spreading pattern of a typical influenza season is relatively well characterized, other patterns may emerge beyond the typical one that contribute to the large variability of influenza spatial transmission observed in epidemiological data.

In the context of influenza epidemics in France, our aim is to identify and classify possible deviations from typical patterns identified in previous work[23]. By studying influenza spatial propagation on a season-by-season basis through a long historical dataset (30 years) we go beyond the description of the robust properties of seasonal waves and build an ontology of possible spreading patterns. In addition to focusing on the peak time, we also aim to consider the patterns that may emerge at the onset of the epidemic [58] and characterize them in a systematic and quantitative way. We classify seasons according to similar onset configurations and similar peak configurations, we then put in relation the two classifications to assess how seasons shift classes from onset to peak, i.e. tracking the potential similarity of flu spreading patterns as the epidemic unfolds throughout the country. Finally, we assess how patterns of seasonal influenza epidemics shift from one class to another depending on demographic, virological, environmental, and mobility drivers. We provide in this way novel insights that were hidden in prior analyses.

## Results

Our analysis is based on influenza-like-illness incidence data for France collected by the French Surveillance Network of general practitioners (Reseau Sentinelles) [58, 10]. We consider weekly time series of incidence rate for the 22 regions of France (NUTS 2 level) for 30 influenza seasons, ranging from 1984-1985 season to 2013-2014 season, and including the H1N1 pandemic season of 2009-2010. For each season we denote the regional onset time as the week of start of the epidemic period at the regional level (see Methods), and the regional peak time as the week with the highest incidence in each region. Regions not experiencing an epidemic period during a season are not considered in the analysis. A large variability is observed across seasons. National epidemic profiles differ substantially in terms of epidemic activity, timing, and duration (Figure 1a). Peak times range from week 49 (month of December, seasons 93-94, 03-04 and 09-10) to week 14 (Month of April, seasons 94-95 and 97-98), with 9 of the seasons peaking before week 2 in January (denoted here as early seasons) vs. 21 peaking after week 6 (late seasons, while instead peaks between week 2 and week 6 are named winter seasons in the following). Attack rates vary considerably, from mild to moderate to severe epidemics. The median time from onset to peak of the epidemic is one month (95% CI [2-7] weeks). Such heterogeneity is largely maintained at the regional level with variations in several indicators (Figure 1b). Almost all regions (median 21 out of 22) experience an epidemic in each season considered. Regions cross the epidemic threshold at different times during the season. If we quantify the spread of the onset time as the time it takes for the percentage of regions reaching the epidemic onset to grow from 10% to 90%, we find it varies from a minimum of 1 week (93-94 season) to a maximum of 9 weeks (85-86 season) (Figure 2a and Supplementary Figure S1). The spread of the regional peak time is comparable, showing a smaller median (3.0 95%CI [1.4-7.9] vs. 3.8 [1.6-8.3]) but still preserving a large variability (Figure 2b).

**Figure 1:**
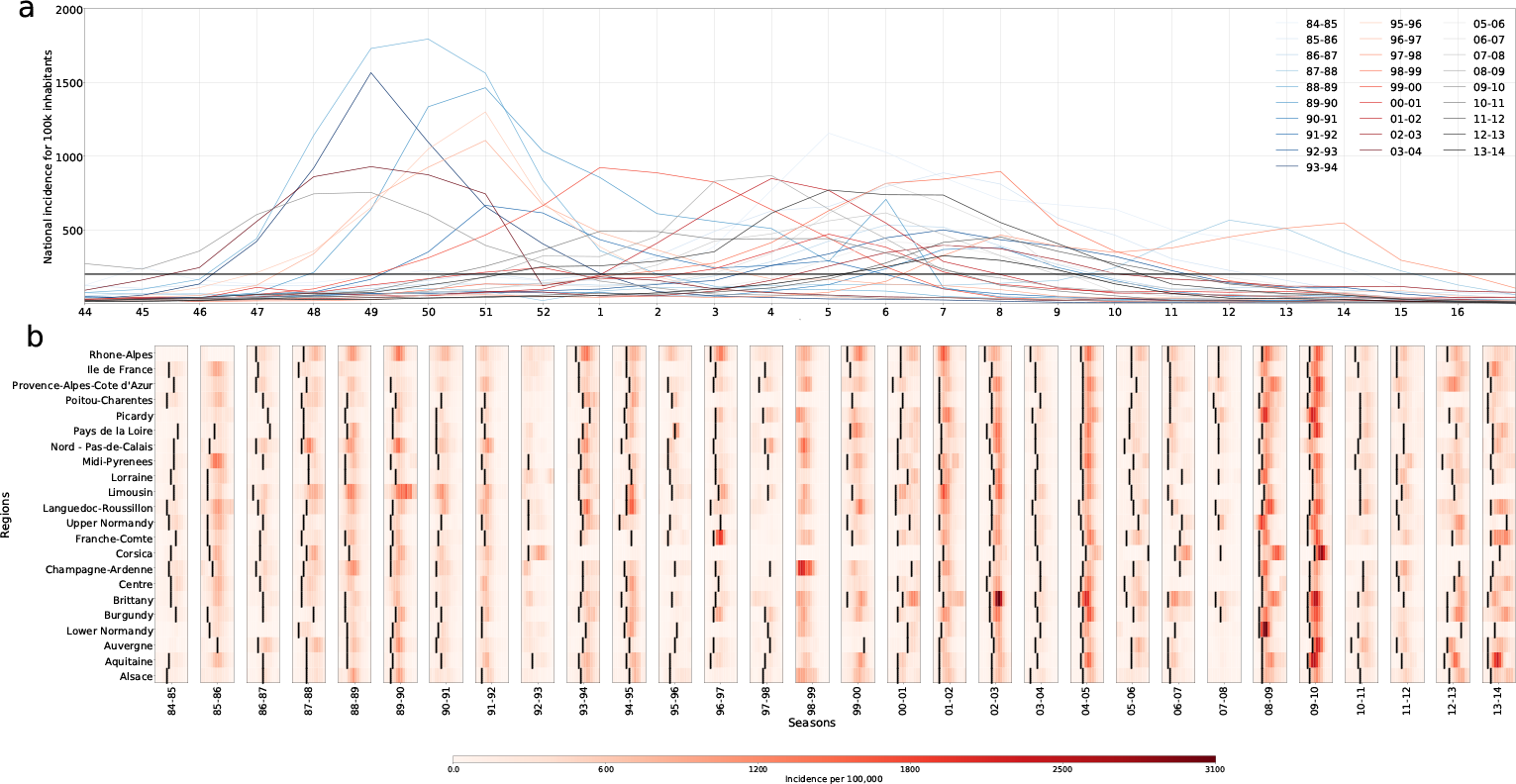
ILI incidence. (a) National ILI incidence for 100,000 inhabitants. (b) Regional ILI incidence for 100,000 inhabitants. The time-interval shown for every season is a 12 week window, centered on the peak of ILI activity (the median value of region’s peak time). Onset time is indicated by the black line.

We now seek to better characterize the heterogeneity observed in the temporal evolution of influenza seasons at regional level. To identify the regional timing patterns for the onset of influenza epidemics, we define for each season s an onset time vector **o**^*s*^, whose element 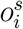 reports the detrended onset time of region *i* in season *s*. Similarly the peak time vector **p**^*s*^ has components 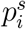 reporting the detrended peak time of region *i* in the season. We then compare different seasons and identify those having similar onset timing patterns by introducing the distance 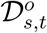 between season *s* and season *t* as:

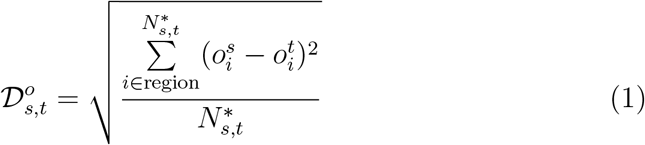

where the sum goes over the 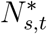 regions experiencing an epidemic in both seasons (regions whose incidence never crosses the epidemic threshold are not considered in the computation). 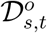 is designed to associate small distances to seasons with similar regional timing pattern, discounting for differences in absolute timing, i.e. early vs. late epidemics. If two seasons share exactly the same onset timing pattern, then 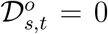. We call this quantity onset distance, and we similarly define the peak distance 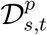 as the analogous quantity computed over the peak time vector **p**^*s*^.

**Figure 2:**
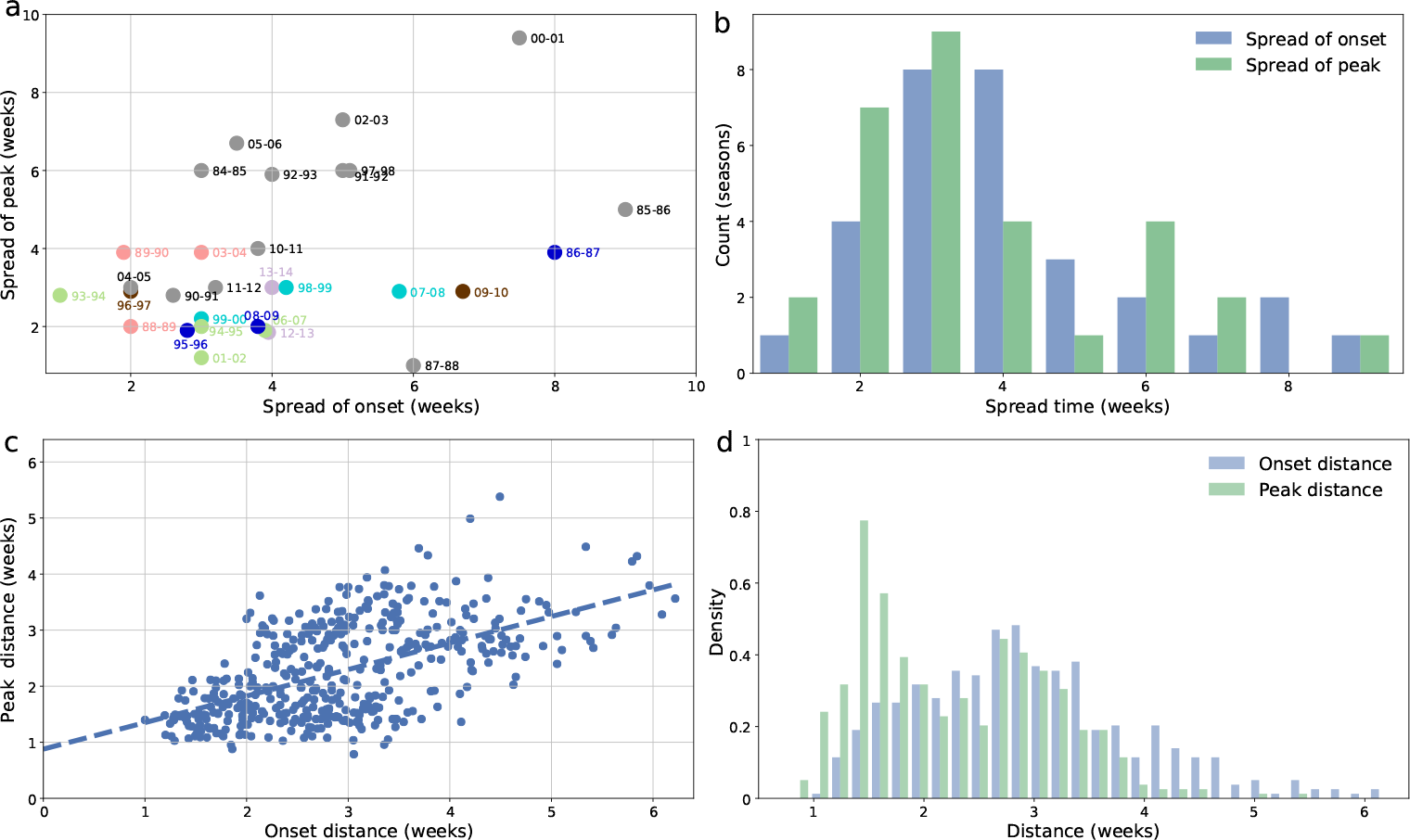
Spread time and distance for both onset and peak. (a) Scattered plot of the spread time of the peak vs. the spread time of the onset, considering all seasons. Seasons are color-coded according to the clustering at the peak (see Figure 3). (b) Spread time distribution for onset and peak expressed in weeks. (c) Scattered plot of peak distance vs. onset distance. Points correspond to all the possible pairs of seasons on which the distance is computed. The dashed line marks the best fit for linear regression. (d) Distribution of onset distance and peak distance values.

Start and peak distance are moderately correlated (*R* = 0.56, *p* < 10^−5^) with *D*^*p*^ displaying a narrower distance, though still highly variable (1.44 – 3.62 weeks, based on the 5th and 95th percentiles) (Figure 2c,d). The peak distance distribution displays two peaks, the highest at around 1.5 weeks and the lowest at around 2.8 weeks, the latter coinciding with the single peak reported for the onset distance distribution.

Next, we classify influenza seasons in terms of their similarity of timing patterns at regional level for the onset and the peak, expressed through the distance *D* computed on the time vectors. We cluster seasons via a complete-linkage agglomerative clustering procedure [30] based on *D*^*o*^, or *D*^*p*^ (Figure 3a). This procedure yields two clustering structures, one capturing similarities between timing patterns of regions at the epidemic onset and one capturing similarities at the epidemic peak (Figure 3b). We then relate these two structures via an alluvial diagram [47] (Figure 3c) mapping the pattern at start with the corresponding pattern at peak, to track how seasons change classification throughout the season.

Out of the 30 seasons under study, we obtain 3 group clusters (O1, O2, O3 in Figure 3c) and 21 single-season clusters (or monoids) for the onset, compared to 6 group clusters (P1, P2, P3, P4, P5, P6 in Figure 3c) and 13 monoids for the peak. Onset group clusters are made of 3 seasons each, whereas the size of the peak group clusters range from 2 to 4 seasons. Most importantly, the alluvial diagram allows us to assess how timing patterns of influenza epidemics change from onset to peak. The analysis shows the emergence of seasons with a similar peak pattern from seasons having a larger distance at the onset of the epidemic. This is observed for seasons 12-13 and 13-14 (grouped in P1) or seasons 98-99, 99-00 and 07-08 (grouped in P2). Also, two of the onset group clusters split when reaching peak time (O1 and O3), whereas 94-95 and 01-02 seasons maintain the similarity of their timing patterns across time (from O2 to P4). All peak monoids but one (11-12 season) are also monoids at the onset of the epidemic. Maps of the group clusters and monoids reveal the strong spatial signatures between seasons of the same cluster (Figure 4 and Supplementary Figure S2). The pattern is given by a large number of regions having the same detrended timing in all seasons of the group cluster (e.g. 8 out of 22 in P1: Bretagne, Pays De La Loire, Haute-Normandie, Centre, Limousin, Île de France, Bourgogne, Franche Comté; 9 in P3: Aquitaine, Poitou-Charentes, Limousin, Languedoc-Roussillon, Rhône-Alpes, Bourgogne, Franche Comté, Champagne-Ardenne, Lorraine), or contributed by several regions at 1 week apart (e.g. 9 regions in P3 or P4 and 11 regions in O2). Focusing on regions with the same detrended timing, we observe that in some clusters they are geographically contiguous (P1, P5, O3), whereas in others they are sparsely distributed (P6). Four regions (Burgundy, Haute-Normandy, Limousin, Île de France) belong to the set of regions with the same detrended timing in at least 50% of the peak clusters (Supplementary Figure S3). Monoids are instead characterized by a large heterogeneity of the timing patterns of regions that strongly differs across seasons (Figure 4c).

**Figure 3:**
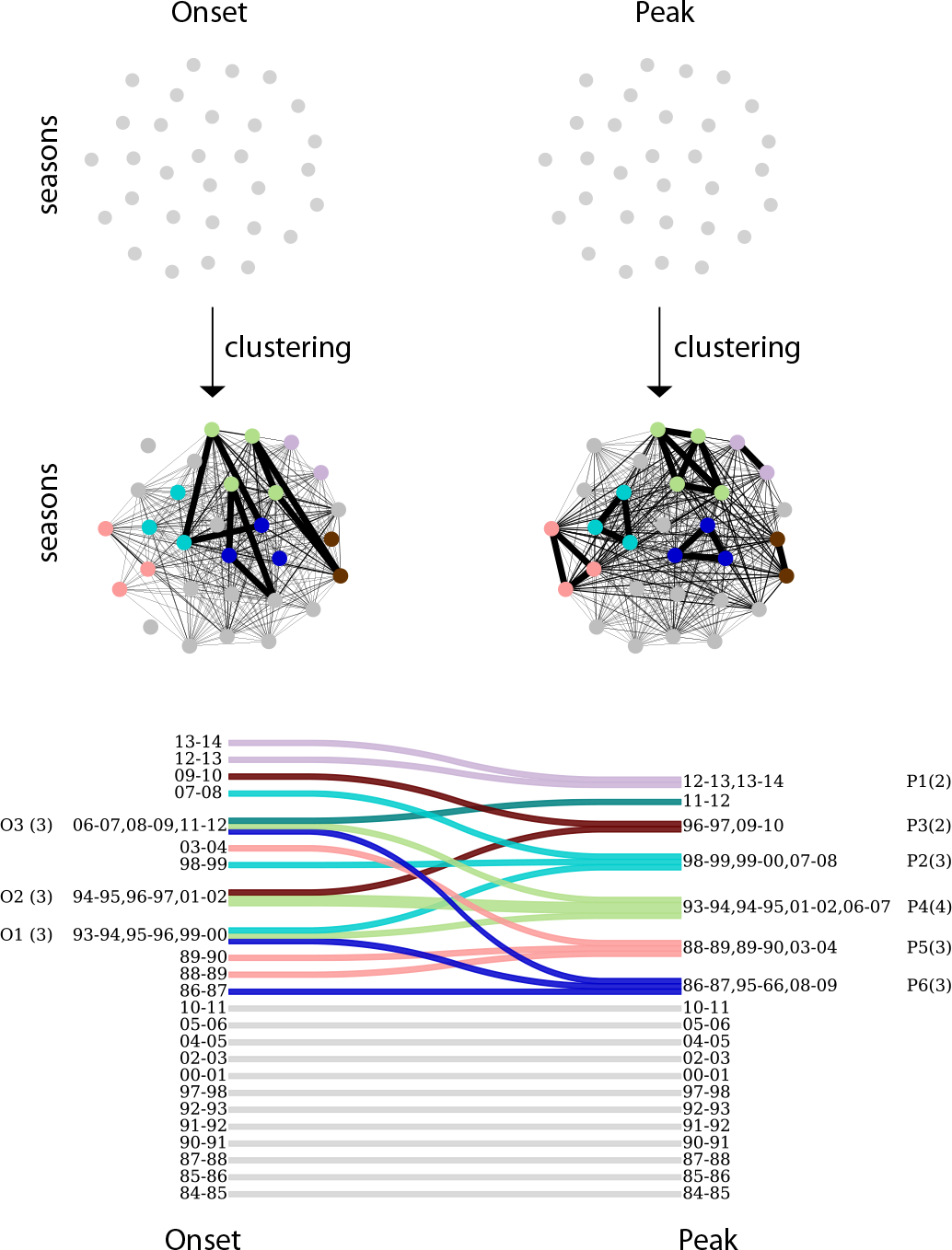
Seasons clustering. (a) Schematic visualization of seasons clustering. Seasons are clustered according to the similarity between their timing patterns, at the onset and at the peak. All seasons in a cluster are within clustering distance from each other. Seasons are color-coded according to the clustering at the peak. (b) Changes in clustering from onset to peak. Onset (left) and peak (right) clustering are shown, with every cluster presented on a different line. The change in timing patterns from onset to peak is represented by mergers/divergences in the lines connecting the two clustering structures. Group-clusters are numbered for future reference, with the number of seasons composing each cluster shown in brackets. Seasons are color-coded according to the clustering at the peak.

We now turn to the analysis of clusters with respect to epidemiological, environmental, and human mobility factors, and focus on the differences between the class of group clusters and the class of monoids. Monoids exhibit large fluctuations in the standard deviation of the regional peak time, with few seasons showing a higher degree of synchronization at the peak (87-88, 90-91, 04-05, 10-11, 11-12) with respect to all other monoid seasons (Figure 5a). Peak group clusters show standard deviations significantly smaller than in monoids only for P2, P4, P5, and P6 clusters (*p* < 0.05). This extends to all group clusters when onset time is considered (*p* < 0.05).

Seasons sharing similar timing patterns at peak are often characterized by higher influenza activity compared to isolated seasons. Four peak group clusters (P2, P3, P5, P6) present larger regional attack rates than monoid seasons (*p* < 0.005), whereas clusters P1 and P4 are not significantly different from monoids (Figure 5b).

Clustered seasons present early epidemics more frequently than expected (*p* < 0.05), and late epidemics less frequently than expected (*p* < 0.05), with monoids showing the opposite behavior (Figure 5c). No difference is observed when considering the onset time. The correlation of the regional ILI time series with the corresponding temperature time series is moderately negative for almost all peak monoids, whereas it can become positive for seasons sharing the same pattern at the peak.

**Figure 4:**
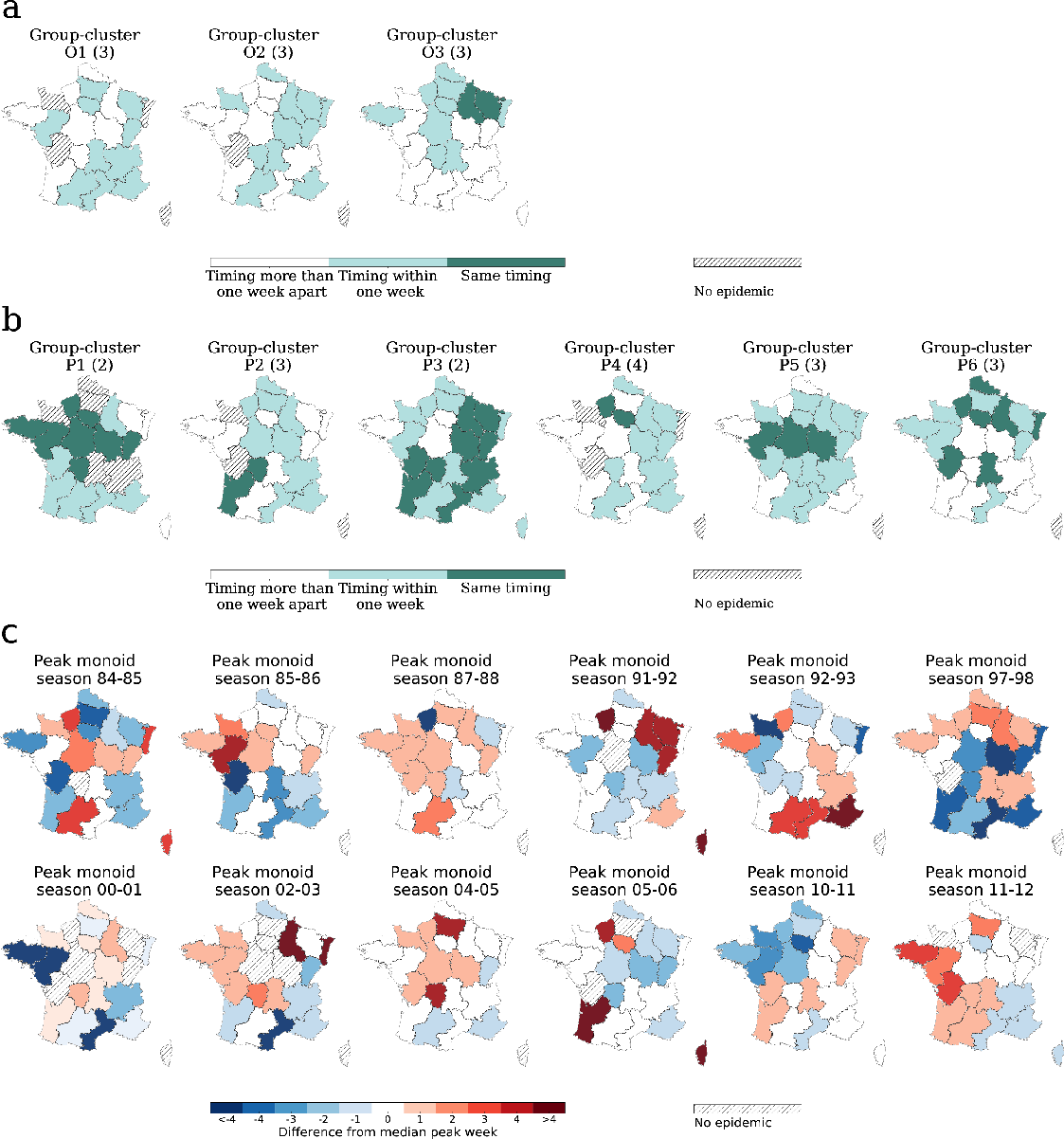
Geographic signature of clusters. (a) Map of France displaying the regional differences in the detrended timing across seasons of the same onset group-cluster. Dark colored regions present the same relative timing in all the seasons, lighter colored regions present a relative timing that is within one week from each other, and white regions present a relative timing that is more than one week apart from each other. Regions having no epidemic in one or more seasons of the clusters are indicated. (b) Map of France displaying the regional differences in the detrended timing across seasons of the same peak group-cluster. The same color-code is used as in panel a). (c) Map of France showing the detrended timing of peak monoids. White regions have a peak time that equals the median value over all regions. Red regions peak later than the median and blue regions before. Regions having no epidemic are indicated. Season 90-91 is not shown, because of the large number of regions not experiencing an epidemic (13 out of 22).

Finally, no association with the dominant strain at the national level is found for the identified clusters.

To better relate the emergence of the regional timing patterns with effects of synchronization at the peak, we compute the pairwise synchronization probability for regions *i* and *j* as the percentage of seasons in which *i* and *j* are synchronized (i.e. their ILI incidence peaks at exactly the same week). We compute this probability considering all seasons under study 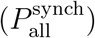, only seasons showing similar patterns at the peak 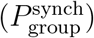, and only isolated seasons in our clustering classification 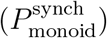. Figure 6 shows an illustrative example of the pairwise synchronization probability for the two most populous regions of France – Île-de-France and Rhône-Alpes. Radial patterns of synchronization appear evident in the group cluster seasons, together with long-range synchronization (e.g. Île-de-France and Midi-Pyrénées, or Rhône-Alpes and Nord-Pas-de-Calais). When considering monoid seasons, radial patterns are far less pronounced (Rhône-Alpes) or largely absent (Île-de-France). Such difference in pattern between seasons clustered in groups or isolated is lost when inspecting the synchronization probability computed on all seasons.

**Table 1:** Correlation values between synchronization probabilities and spatial/anthropological factors (Mantel test, n.s. stands for “not significant”).

| | $P_{\text{monoid}}^{\text{synch}}$ | $P_{\text{group}}^{\text{synch}}$ | $P_{\text{all}}^{\text{synch}}$ |
| --- | --- | --- | --- |
| Distance | -0.225** | -0.384** | -0.442** |
| Product of population | n.s. | n.s. | n.s. |
| Flight passengers | n.s. | n.s. | n.s. |
| Commuters | n.s. | 0.195* | 0.182* |

\* $p < 0.05$
\*\* $p < 0.01$

To quantitatively measure this effect and assess its possible drivers, we measure the association between the probability of synchronization and several demographic and mobility indicators via a Mantel test (Table 1). Pairwise synchrony significantly decreases with distance in both types of clusters (mild correlation) and over all seasons (moderate correlation). No association is found with the product of the population sizes of the donor and recipient regions or with air travel fluxes across all groups of seasons considered. Commuter flows instead display a weak positive correlation with the probability of synchronization considering all seasons and group-clustered seasons, whereas no association is found for monoids.

**Figure 5:**
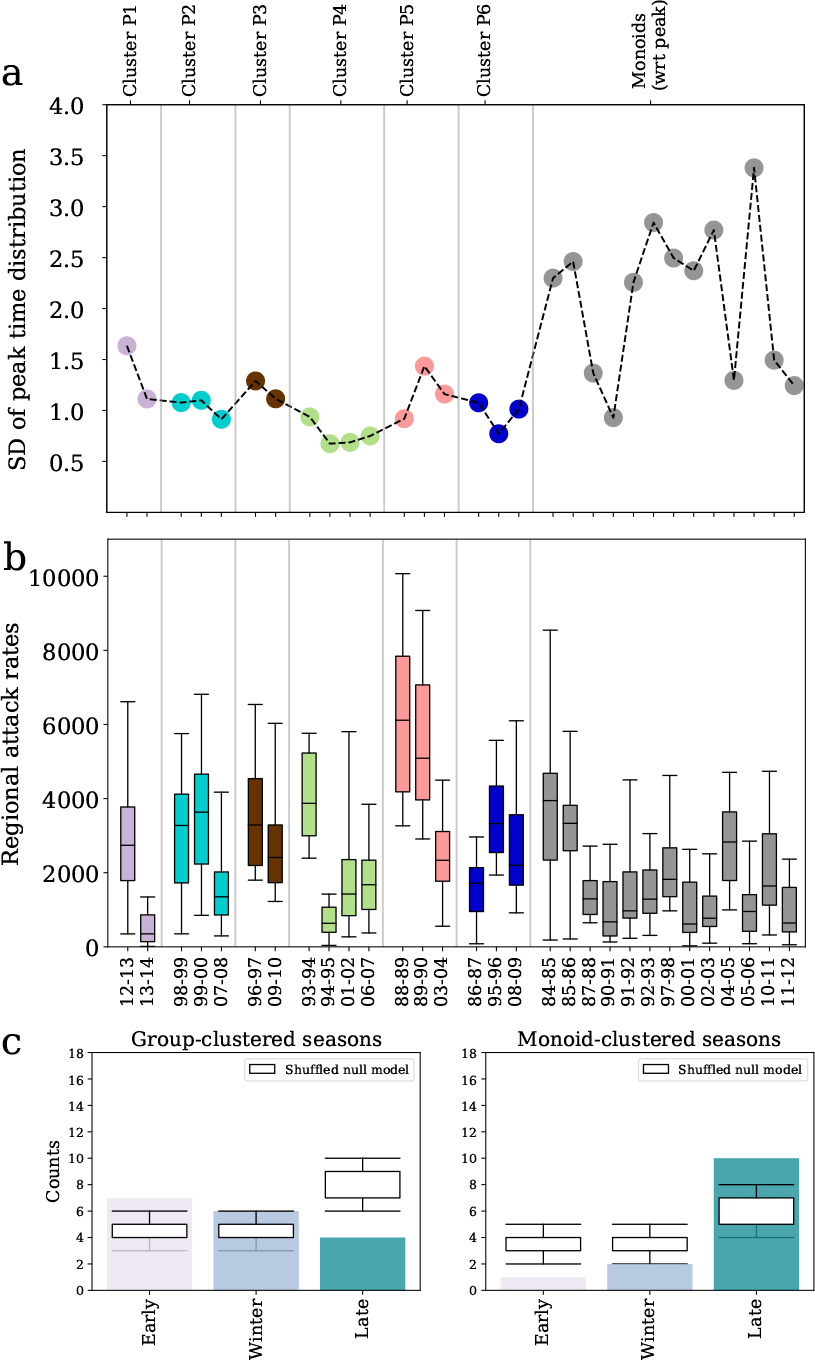
Synthetic indicators characterizing seasons clusters. Seasons that form group-clusters with respect to peak time are highlighted with the corresponding cluster colour (see Fig. 3), while seasons that are monoid-clustered are shown in gray. (a) Standard error of the distribution of regional peak time by season. (b) Box plot of regional attack rates by season. (c) Seasons classification with respect to absolute peak time. Number of seasons that have an early, winter or late peak time (see main text) for group-clustered seasons (left), and for monoid-clustered seasons (right) with respect to peak time. The classification is compared to a random null model (boxplot).

**Figure 6:**
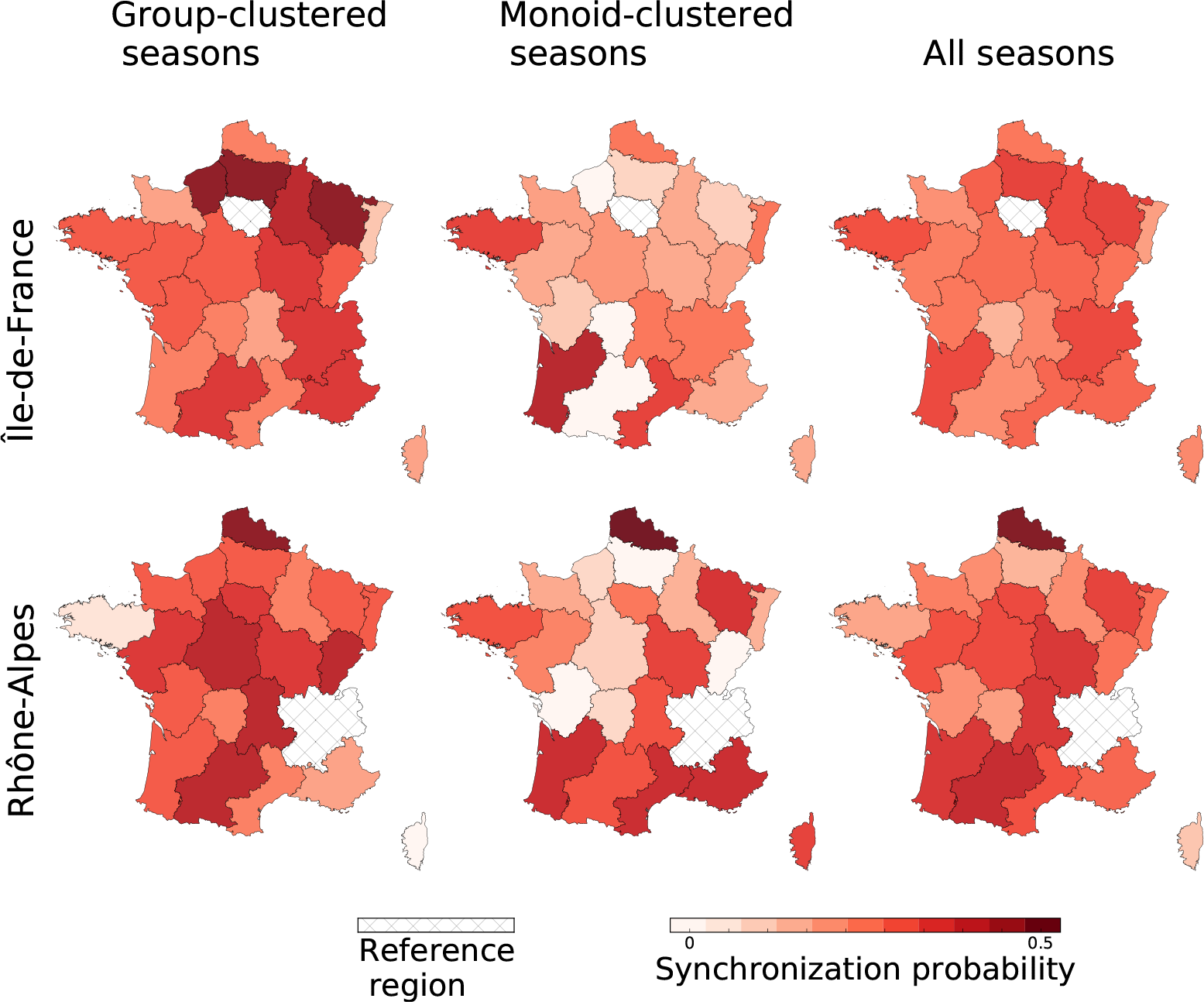
Synchronization probability. Color-coded pairwise synchronization probabilities 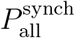, 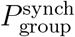 and 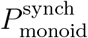 of all regions with Île-de-France (top row) and Rhône-Alpes (bottom row).

## Discussion

Elucidating the spatiotemporal patterns of the spread of influenza epidemics is of great importance for preparedness and control. Here we study 30 influenza seasons in France at the regional level and introduce a novel method to characterize the variability of spatial transmission of influenza. We consider a clustering approach to reduce the observed variability by identifying a limited number of regional configurations at the epidemic onset (onset timing patterns) and at peak time (peak timing patterns) shared by multiple seasons. We characterize their properties, and assess how seasons shift between classes of patterns depending on epidemiological, environmental, demographic and mobility factors.

The cluster analysis yields two important findings. First, two relevant classes of clusters emerge: group clusters composed of seasons sharing recurrent diffusion patterns and monoid clusters made by single seasons whose patterns strongly differ from all others. This is observed both at the onset and at the peak of the epidemic, although the mechanisms for pattern formation at the two stages of the epidemic are different and are not trivially captured by epidemiological or virological signatures at the national level (e.g. intensity of the epidemic activity, or dominant virus type). Second, the current knowledge characterizing a typical influenza epidemic only applies to a subset of seasons, specifically those exhibiting recurrent patterns, whereas unique patterns, though representing roughly 50% of the seasons, show different properties.

A large number of works focused on the characterization of influenza spatial propagation in different countries and at different geographic scales [31, 23, 65, 49, 22, 38]. The novelty of our approach is to systematically identify recurrent patterns across seasons, distinguish them from unique patterns in a fully hierarchical way, and to connect the patterns at epidemic onset with those at peak time. This is made possible by the long timeframe under study. We find that as influenza incidence increases from the epidemic threshold to the peak, patterns become more similar thus presenting a stronger clustering at peak compared to patterns at the onset. Highly contagious viruses [61, 32] and strong population coupling are generally associated with faster and more homogeneous spatial spread [46, 33, 12, 28]. This is further increased by the small scale of the country, as reported in the case of Israel displaying a high synchronization across cities [32]. Our study however shows that neither the standard deviation of the peak time distribution across regions nor the regional attack rate are able to discriminate in a definitive way between recurrent and unique patterns. While synchronization is indeed a mechanism to group seasons in the same cluster, similarity can be achieved also in absence of a high synchronization (i.e. homogeneity of timing) as long as the regional heterogeneity is close across seasons. In addition, we find clusters that are built on a large number of regions whose timing differs of one week only (P2, P4), thus maintaining low values of *D*^*p*^. Notably, Île de France plays this role in the majority of the peak clusters.

The spatial pattern of influenza transmission in France is predominantly localized and driven by the distance between regions, with a stronger effect observed on recurrent patterns compared to unique ones. This is in line with previous observations of wave-like spreading behavior [61] and of strong correlation between neighboring regions reported for a typical influenza season in the country [23]. Similar results connecting distance and synchronization were also obtained for the US through wavelet analysis [61], spearman rank correlation [56] or statistical modeling [22]. The effective role of distance in the spatial transmission of influenza has been put in relation with different modes of mobility of individuals. Commuting has been identified as an important driver for short-range dissemination [21], though the narrow nature of its range of connectivity is not able to explain alone the broader spatial propagation of influenza [22]. For France we find a positive, though weak, correlation considering all 30 seasons, signaling that a combination of different modes of travel may compete at the scale of the country [12]. No support is indeed found for air travel, in line with previous works on influenza in France [21, 23], likely because the majority of internal mobility relies on ground transportation. On the global scale, air travel is instead associated to the large-scale propagation of seasonal influenza epidemics [48, 36, 64], and is known to be an important driver for the dissemination of emerging infectious diseases [17, 45, 44], including influenza pandemics [13, 34, 27]. At the national level, results do not appear to be conclusive [61, 22, 18, 12, 16] and seem to depend on the scale of the country, with smaller countries like for example France [23] and Israel [32] clearly excluding a dominant role of mobility by air for influenza spread compared to the US [18, 20, 22].

Our clustering approach shows however that these spatial transmission properties are specific only to a subset of influenza epidemics, namely those exhibiting recurrent patterns. The dependence on commuting observed when all seasons are considered is recovered for group clusters but is absent in seasons with unique patterns. For the latter, the dependence on the distance is also weaker, thus signaling that, besides an underlying weak localization, the mode of spread is less marked spatially, as evident from the maps of Figure 4 and Figure 6. No dependence on mobility and demography is found, so other factors must be at play that concur to the emergence of these unique patterns.

We find a clear dependence on the temperature only for monoids, exhibiting a negative correlation with temperature profile [41] (Supplementary Figure S4 in the SI). This is likely induced by the unexpectedly large number of late epidemics, for which the ILI peak follows in time the week of minimum temperature. Temperature profiles for France usually have their minimum in between the end of January and the first week of March (5th and 95th percentile, respectively) while ILI activity has a greater variability in peak time, ranging from the beginning of December to mid March (5th and 95th percentile, respectively).

Previous work has highlighted that different types/subtypes of influenza present different circulation patterns [61, 48, 14]. Season classification in terms of dominant strain at the national level however does not allow us to establish a connection between virology and spreading patterns. Possible causes may include the small size of the sample, given that the national strain dominance (defined as >50% of isolated samples) is available for 25 seasons (6 B seasons, 8 H1 seasons, and 11 H3 seasons) and is compared to a relatively small number of group clusters. In addition, co-dominance of different influenza strains or co-circulation of influenza virus with other respiratory pathogens (as e.g. respiratory syncytial virus [51, 59]) are expected to possibly lead to non-trivial interactions and likely a mixing of different spatial patterns within the same season. Finally, our results seem to suggest that regional heterogeneity in strain dominance is taking place in the country, with one strain dominating at the national level while different configurations of sub-dominant strains compete at the local level. For example, seasons 08-09 and 11-12 are clustered together at the start, but not at the peak. Though sharing the same dominant strain at the national level (H3), season 08-09 displays influenza B becoming dominant in some regions throughout the season, whereas in season 11-12 type B is never dominant [6, 5]. Spatial heterogeneity in influenza strain dominance is often reported at a larger scale across countries in Europe [3, 2], with countries exhibiting different strains dominating during the same influenza season. The underlying mechanisms are however still poorly understood. Virological data with regional resolution would be needed to further investigate this aspect in France, however these are available starting 2014-2015 season only, following a change in surveillance protocols.

Clustering is less structured at the onset compared to the peak and displays a larger number of unique patterns. This may be related to varying configurations of case importations from abroad that change with seasons, and it is indeed in line with the large variations reported for the estimates for the external seeding parameter of the model fitting influenza diffusion in the US [22].

While we are able to discriminate between recurrent patterns that confirm previous knowledge on typical influenza spatial transmission and unique patterns exhibiting different properties, some aspects emerging from our classification still elude our quantitative interpretation. For example, we find some seasons shifting from similar onset patterns to different peak patterns (e.g. seasons 93-94,95-96 and 99-00). The demographic, virological, and environmental factors we tested are not able to explain the shift. Also, mobility is unlikely to be a possible driving factor as the shift would require a marked topological change in commuting flows over time that is not reported by national statistics. We expect possible factors therefore to be dependent on space and to change across years. One such factor could be a different virological signature at the local level, as discussed before. A second possible cause could be a spatially different vulnerability of the population to the disease, due to different age structures (e.g. the fraction of individuals under 20 years of age varies from 21% in Corsica to 27% in Nord-Pas-de-Calais, data of 2015 [9]) or varying vaccination rates per region. A statistical model fitting ILI incidence data in the US found a large variation in the parameter value fitting population susceptibility to influenza across seasons [22]. This seems to support our hypothesis on the role of different profiles of regional immunity that may change in time. We repeated the analysis on ILI age structured data and obtained similar results, suggesting that we are unable to uncover these mechanisms on surveillance data only. Validation on serological or vaccination coverage data at the regional level is prevented by the lack of exhaustive data, but presents an interesting avenue to explore in future data collection developments.

Some limitations affect our study. First, we use ILI data as a proxy for the spatio-temporal evolution of influenza epidemics. This is in line with a large number of previous works [62, 61, 23, 32, 29, 22], as ILI data are known to be a reliable proxy for influenza incidence when activity is sustained [63, 42, 60]. Close to the threshold the agreement is expected to be lower, thus possibly affecting the identification of the onset time of the epidemic especially in smaller regions. For this reason, we also tested different regional epidemic thresholds finding that our results are robust against this change. Second, different definitions of the onset times may be proposed [22]. Here we decided to adopt the national definition of the epidemic period remapped to the regional scale. Third, the surveillance network of sentinel general practitioners is not a representative sample distributed in the country. Recent work proposed different techniques to correct ILI data in France against this possible bias [54, 55]. We checked that these corrections would leave the peak time invariant and would lead to a small change of the threshold value that would however not alter the identification of the onset time, as discussed above. Fourth, our classification depends on the clustering threshold chosen. While we tested the robustness of our choice in the sensitivity analysis (see Supplementary Figure S5 in the SI), our approach maintains an arbitrary degree in the definition of the similarity across seasons that may be tuned according to the study objective. Finally, we note that starting 2016 France adopted a different administrative subdivision in 13 regions following an aggregation of the 22 regions considered here. Our study could be easily extended in future work as surveillance data are currently collected by keeping both subdivisions. Using the 22 regions subdivision would allow us to maintain the comparison with past seasons and also to have a higher granularity to better appreciate spatial patterns.

## Conclusions

Our study introduces a novel method to better classify different spatiotemporal trends in seasonal influenza activity. A larger variability is observed at epidemic onset with respect to peak time. The clustering at peak time reveals a more pronounced spatial diffusion for seasons exhibiting recurrent patterns compared to seasons characterized by unique patterns. Commuting plays a measurable role in shaping recurrent influenza spatial patterns whereas it is found to play no role in unique patterns seasons. This provides new insights that were hidden in prior analyses that were not capable of assessing the full spectrum of possible configurations. Groups of seasons shifting from similar onset patterns to different peak patterns do not find a definitive explanation in the present study but open the path to future perspectives assessing viral strain dominance and population immunity at a higher resolution than what currently available.

This work is the first systematic classification of influenza seasons in terms of diffusion patterns at onset and at peak, but the methodology described here is completely general. Applying this method to countries of different scale within the same study period may help identify season-specific properties that are robust in the classification and shed light on the interplay between geographical scale and observed propagation. Also, the comparison between clustering of influenza seasons and of other respiratory disease epidemics may yield novel insights into the relation between pathogens and associated spreading patterns.

## Material and Methods Data

### Data

ILI incidence data is provided by the French GP surveillance network adopting the following ILI definition: a sudden fever of over 39°C with myalgia and respiratory symptoms[25]. Incidence curves are smoothed with a 3-week moving average to avoid noisy fluctuations. Incidence data at the regional level is considered for all age classes aggregated, and also broken down for children (less than 20 years) and adults (otherwise) classes for age-specific analyses.

Mobility data are obtained from two sources. Commuter flows between different departments (NUTS 3 level) are extracted from data of the French National Institute of Statistics and Economic Studies (Insee) for 2011 [8] and aggregated at the regional level (NUTS 2 level). Air travel fluxes between French regions are extracted from the IATA worldwide airport database [7] aggregating all airports in the same region. A total of 60 commercial airports exist in France with 193 within-country connections.

Temperature data are obtained from the European Climate Assessment & Dataset (ECA&D)[35, 4]. Daily data from all weather stations in a given region are aggregated to compute an average weekly temperature for the region. Data extended up to 2005, and for some regions no temperature data was available.

### Epidemic threshold and onset time

For every season, the regional onset time of the epidemic is defined as the first of two consecutive weeks for which the ILI incidence rate is above 150 cases for 100,000 inhabitants. No standard definition is available for the onset of the epidemic period at the regional level, and currently the Sentinel Surveillance System adopts an approach similar to the one used at the national level. By comparing ILI data [10] and virological data [1] for the same season in France, the above threshold value is obtained by imposing that with 95% confidence the percentage of positive swabs is at least 15% of the total ILI samples (Supplementary Figure S6). We also tested different values for sensitivity. If the incidence rate is never above the threshold value, the region is considered as not experiencing an epidemic and is excluded from the analysis.

## Author’s contributions

PC, CP and VC conceived and designed the analysis. PC performed the analysis. PC drafted a first version of the manuscript. PC and VC wrote the final version of the manuscript. All authors contributed to the interpretation of the results, the writing of the manuscript, read and approved the final manuscript.

## Acknowledgements

We thank Shweta Bansal for fruitful discussions on this study.

